# Parameter tuning facilitates the evolution of diverse tunneling patterns in termites

**DOI:** 10.1101/836346

**Authors:** Nobuaki Mizumoto, Paul M. Bardunias, Stephen C. Pratt

**Affiliations:** School of Life Sciences, Arizona State University, Tempe, AZ, USA 85287; Department of Biological Sciences, Florida Atlantic University, Boca Raton, FL 33431

## Abstract

The nest structures built by social insects are complex group-level patterns that emerge from interactions among individuals following simple behavioral rules. The theory of complex systems predicts that there is no simple one-to-one relationship between variations in collective patterns and variation in individual behaviors; therefore, it is essential to know how actual behavior evolves to change pattern formation. Here we demonstrate that the evolutionary divergence of termite tunneling patterns is achieved by quantitative tuning of shared behavioral rules, rather than the acquisition of novel behaviors. We compared tunnel formation between two closely related species, *Reticulitermes tibialis* and *Heterotermes aureus*, and found that *H. aureus* builds more highly branched tunnels than *R. tibialis.* Our behavioral analysis and data-based modeling revealed that these species share the same behavioral repertoire, but a quantitative difference in the probability of sidewall excavation leads to diverse tunneling patterns. In contrast, we also found that *Paraneotermes simplicicornis*, which evolved tunneling independently, possesses a distinct behavioral repertoire, but shows convergence of branching patterns with *R. tibialis*. These results elucidate the complex relationship between individual behavior and group-level patterns; in some cases, distinct behavioral rules can produce similar group-level patterns, but in others, a common rule set can yield distinct patterns via parameter tuning. The evolutionary process of collective behavior is flexible and much more complex than we can infer from group-level patterns alone.

## Introduction

The coordinated behavior of group-living animals often creates complex group-level patterns [1]. Among these, nest structures built by social insects play an important role in their ecological success by providing shelter and favorable microenvironments [2,3]. A wide variety of structures has evolved, adapted to each species’ typical environment [4,5]. This leads to the fundamental question of what is the behavioral mechanism underlying the evolution of diverse nest structures? In collective building, group-level structures emerge from local interactions among individuals following simple behavioral rules [1,6]. Thus, different collective outcomes may be obtained either by differentiated behavioral rules or by regulation of a common set of rules to modify the interactions [7]. Theoretical studies have supported the latter model; they predict that diverse nest structures can be explained by parameter tuning, which is the quantitative modification of a single set of behavioral rules shared among species, in ants [8,9], termites [10–12] and paper wasps [13,14]. However, because of the lack of comparative studies, there is no empirical evidence for the sharing of behavioral rules across species, and thus the key factor creating interspecific variation in patterns remains unknown.

Because evolutionary changes of nest structures occur through the modification of building processes, it is essential to clarify the relationship between individual behaviors and group-level patterns. Evolutionary developmental biology has treated similar challenges by elucidating the mechanistic relationship between individual development and phenotypic change during evolution [15,16]. Studies on the molecular basis for divergence and convergence of individual morphology suggest that all scenarios are possible, where different forms may evolve by similar mechanisms [17] and similar forms may evolve by different mechanisms [16]. Applying this perspective to nest structures, which can be seen as the external morphology of a colony [18], social insects may not only use parameter tuning mechanisms but also acquire differentiated behavioral rules during the evolution of collective building.

In this study, we analyze the relationship between individual behavior and collective pattern in the tunneling behavior of termites. Several termite species build tunnels to protect foragers from desiccation and predators [19]. Termite species can vary in tunneling patterns, reflecting species-specific foraging strategies and differences in the distribution of wood resources experienced by each species [20–22]. Moreover, our phylogenetic analysis indicates that tunneling behavior has evolved several times independently in termites (Figs. 1, S1). This provides an opportunity to explore the extent to which behavioral rules are shared among species with different degrees of relatedness.

**Figure 1.**
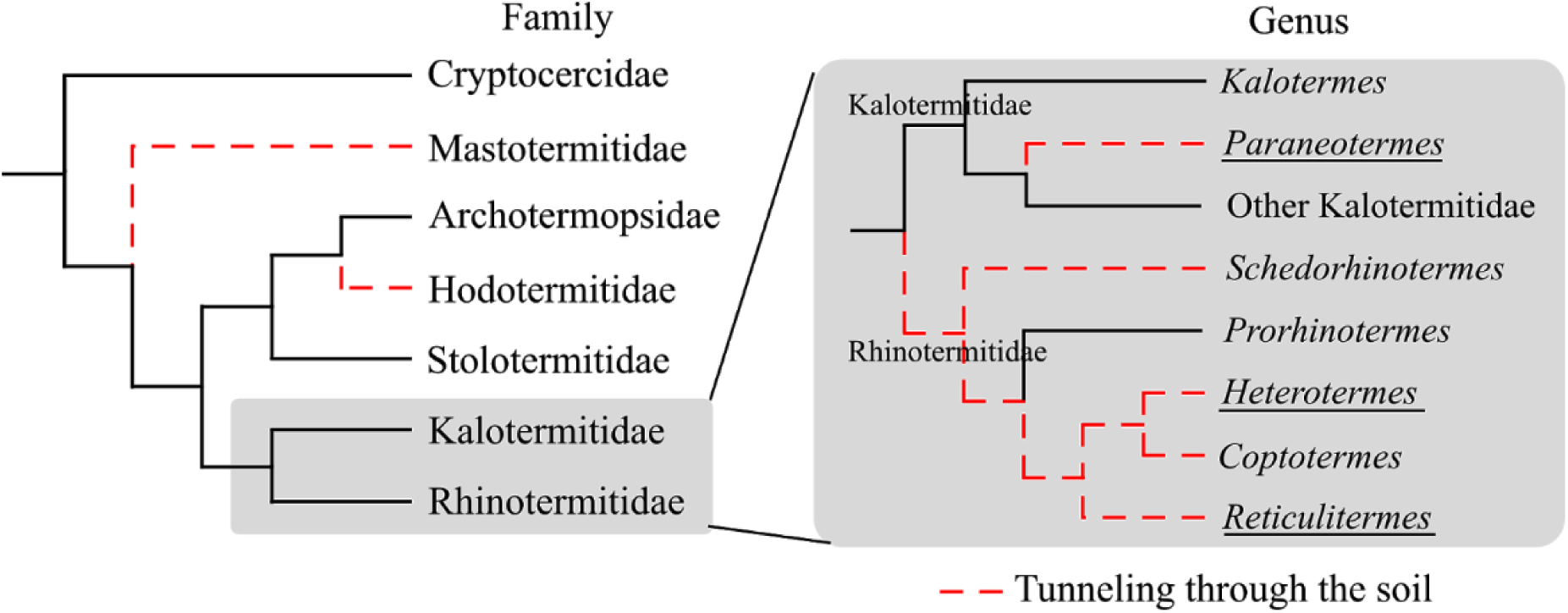
Simplified phylogeny of lower termites (modified from [23,24]) with information on tunneling behavior. Ancestral states were reconstructed with maximum parsimony (detailed in SI text and Fig. S1). Tunneling through the soil has evolved four times independently in Mastotermitidae, Hodotermitidae, *Paraneotermes*, and Rhinotermitidae. In this study, we used three species from the three underlined genera, *Paraneotermes* (Kalotermitidae), *Heterotermes*, and *Reticulitermes* (Rhinotermitidae).

To trace the evolutionary changes of tunneling patterns and behaviors, we used three subterranean termite species. *Paraneotermes simplicicornis* (Kalotermitidae) evolved tunneling independently from *Reticulitermes tibialis* and *Heterotermes aureus* (Rhinotermitidae) (Figs. 1, S1). We observed tunnel development at two different scales: the patterns of tunnel branching and the behavior of each termite. We empirically demonstrate that there is no simple one-to-one relationship between individual behaviors and group-level patterns. We find that *R. tibialis* and *H. aureus* build tunnels with distinct branching patterns from each other by using shared behavioral repertoires, while *P. simplicicornis* builds similar tunneling patterns to *R. tibialis* using a distinct behavioral repertoire. We combine empirical observations and data-based simulations to show that different branching patterns between *R. tibialis* and *H. aureus* result from parameter tuning of the same behavioral rules. Thus, quantitative modification of shared behaviors can play an important role in the evolution of diverse group-level patterns, without any change in the behavioral rule itself.

## Results

### Tunneling patterns

Our observation of tunnel development in a two-dimensional arena detected both convergence and divergence of termite tunneling patterns (Fig. 2A). The most striking feature of group-level patterns is the number of branches. When we counted the number of tunnel faces, we found that *H. aureus* built a significantly larger number of branches than *P. simplicicornis* and *R. tibialis*, while there was no significant difference between *P. simplicicornis* and *R. tibialis* (generalized linear model [GLM]; likelihood-ratio test, χ^2^_2_ = 11.568, *P* = 0.00308; Tukey Contrasts: *R. tib.* - *P. sim.*: *z* = 0.376, *P* = 0.925, *H. aur.* - *P. sim.*: *z* = 3.007, *P* = 0.0073, *H. aur.* - *P. sim.*: *z* = 2.611, *P* = 0.0245; Fig. 2B). Branching events were mainly found near the tunnel entrance (Fig. 2A), where the length of the tunnel before the branches is much shorter than that after the branches (Fig. 2C).

**Figure 2.**
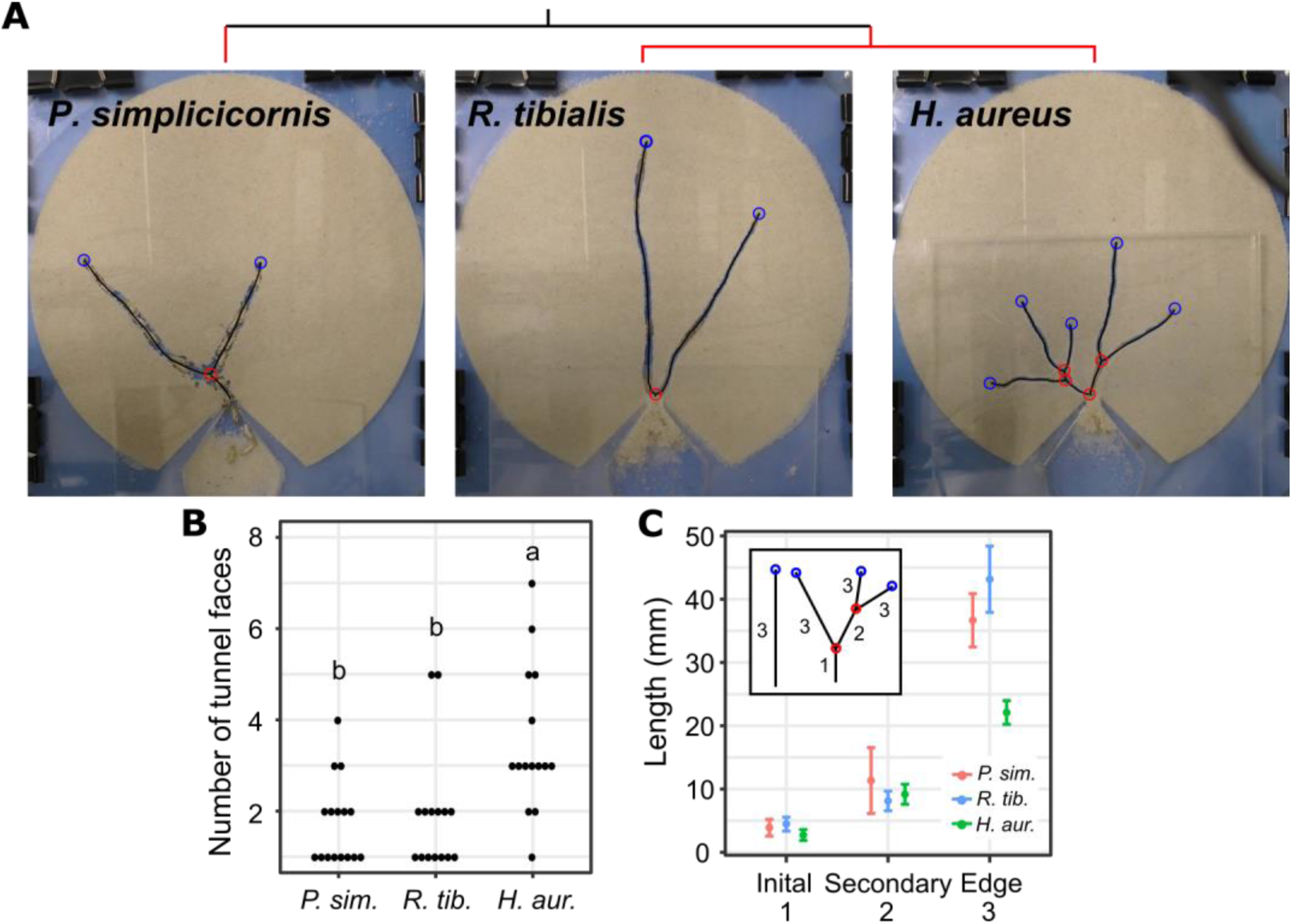
Interspecific comparison of termite tunneling patterns. (A) Typical tunneling patterns of each species. The red lines in the simplified phylogeny above the photos indicate the independent evolution of tunneling. Red circles indicate branching points and blue circles indicate the faces of the tunnels. (B) Comparison of the number of tunnel faces among species. Different letters indicate significant differences (*P* < 0.05). (C) Comparison of the tunnel length when divided into segments. Initial tunnel is a segment from the entrance to the first branch. Secondary tunnel is a segment between two branches. Edge tunnel is the segment reaching the faces of the tunnels. When a tunnel has no branch, it only contains an edge tunnel.

### Individual digging behavior

Individual behavior during tunneling did not correspond directly to the group-level patterns; instead, *P. simplicicornis* used a distinct transporting behavior unlike that of either *R. tibialis* or *H. aureus*. In this behavior, each individual of *P. simplicicornis* excavated sand with its mandibles, formed the sand into a ball with its legs, and then kicked the ball backward to the individual behind it (Fig. 3A; Video S1). This behavior was observed only in this species (Fisher’s exact test, *P* < 0.0001; Fig. 3B) and contrasts with the well-recognized behavior of rhinotermitid termites; these species excavate and carry sand particles using their mandibles (Fig. 3A; Video S2). These different behaviors were associated with different tactics for removing the sand; *R. tibialis* and *H. aureus* individually carry sand out of the tunnel, while *P. simplicicornis* instead forms a bucket brigade of multiple individuals. This was apparent in the fact that *R. tibialis* and *H. aureus*, but not *P. simplicicornis*, finish transportation by compressing clumps of sand particles against the sidewall (Fisher’s exact test, *P* < 0.0001; Fig. 3B). *P. simplicicornis* just kicked sand balls into the tunnel passage, where they were taken over by another individual. In addition, the order of individuals inside a tunnel was maintained in *P. simplicicornis*, where the individual in the 1st-row at the tunnel face was less likely to move back and change positions with 2nd-row individuals (one-way analysis of variance [ANOVA], *F*_2_ = 24.307, *P* < 0.0001; Fig. 3C). Moreover, *P. simplicicornis* visited the tunnel face fewer times (ANOVA, *F*_2_ = 24.892, *P* < 0.0001; Fig. 3C), indicating that they transport a large amount of sand at once using the bucket brigade. Finally, after visiting the tunnel face, 1st-row individuals of *P. simplicicornis* only moved back a short distance as another individual can take over the sand ball. This trend was prominent when the tunnel became longer, where *P. simplicicornis* typically moved back only about 10 mm regardless of the length of the tunnels, while *R. tibialis* and *H. aureus* moved back increasingly long distances as the tunnel lengthened (Fig. 3D).

**Figure 3.**
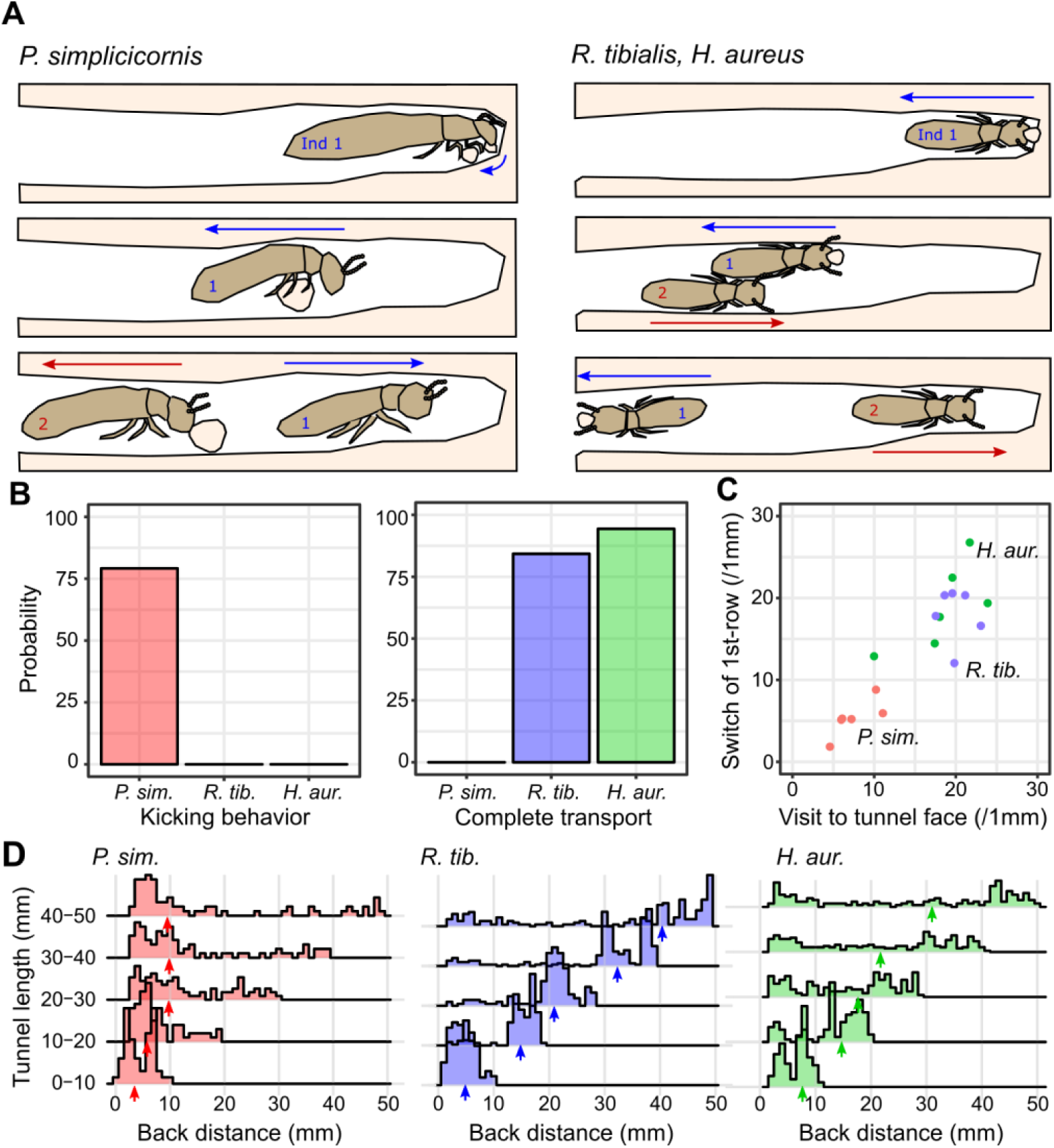
Comparison of individual behaviors during tunneling. (A) Illustration of sand excavation behavior. After excavating sand at the tunnel face, *P. simplicicornis* use their legs to form a ball of sand particles and kick it back behind, where it is taken over by the 2nd-row individual. *R. tibialis* and *H. aureus* instead carry sand particles with their mandibles, and the individual in the 1st-row often changes. (B) Interspecific comparison of the probability to show kicking behavior and completing transportation. Completion of transportation is defined as compressing or attaching sand particles, which are then not used by other individuals. (C) Interspecific variation of the number of visits to the tunnel face and the number of position switches between 1st-row and 2nd-row individuals. (D) Comparison of the distance of backward movements after visiting the tunnel face. Arrows indicate median values.

We further investigated interactions among individuals to specify individual-level differences that might account for the different tunnel patterns of *R. tibialis* and *H. aureus*. During excavation, direct interaction between termites can occur in a clogged tunnel. We observed the behavior of 2nd-row individuals when they found themselves immediately behind the 1st-row individuals who are excavating at the tunnel face (Fig. 4A). We found that *R. tibialis* and *H. aureus* chose from the same behavioral repertoire, either excavating the sidewall or waiting until the 1st-row individual has finished excavation. However, the two species differed in the frequency of these behaviors (Fisher’s exact test, *P* = 0.0035, Fig. 4A). The 2nd-row individuals in *R. tibialis* more often waited, while those in *H. aureus* had a higher probability of beginning to excavate the sidewall (Fig. 4A). In contrast, *P. simplicicornis* showed a distinct behavior not present in the other two species, where the 2nd-row individual took over the transportation of the sand ball kicked back by the 1st-row individual (Fig. 3A, 4A). Thus, phylogenetically distinct species have different behavioral repertoires, while closely related species share the same repertoire but quantitatively modify their use of it, indicating the existence of parameter tuning.

**Figure 4.**
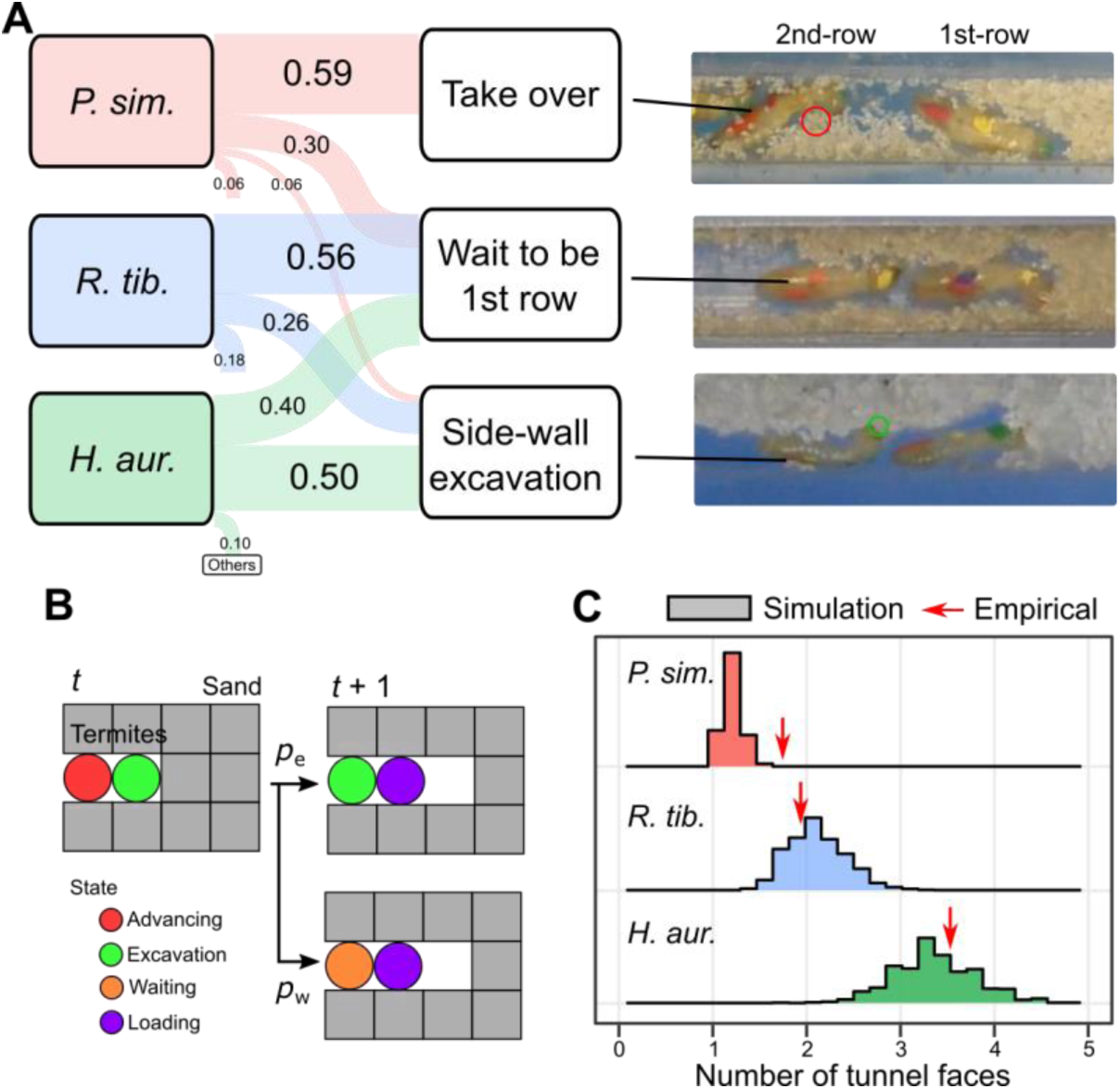
Parameter tuning mechanism for interspecific variation of tunneling patterns. (A) Interaction rules between individuals within a clogged tunnel. In *P. simplicicornis*, a 2nd-row individual often takes over the sand ball the 1st-row individual has kicked back. On the other hand, *R. tibialis* and *H. aureus* wait or excavate side wall, where these two species are different in the frequency of these behaviors. (B) Behavioral rules governed the simulated interactions for *R. tibialis* and *H. aureus*. If a termite finds an excavating individual in front of it, it will choose an action of wait or excavate side wall depending on the probability obtained in the experiments. (C) Comparison of the results of empirical experiments and simulations. Histograms indicate the mean values for every 15 or 16 simulations (N = 1000), while red arrows indicate the mean value of empirical experiments.

### Simulations

We hypothesized that the observed quantitative difference in sidewall excavation between *R. tibialis* and *H. aureus* is the mechanism of branching pattern variation. In fact, it has been reported that such sidewall excavation widens the tunnel and can eventually result in a new branch [25]. To test our hypothesis, we developed a cellular automaton model simplifying the tunneling process (Fig. S2). In the model, each termite moves towards the end of the tunnel from the installation area as long as the space in front is empty. When a termite reaches the face of the tunnel, it excavates the cell’s contents, making it empty. After excavation, the termite moves back some distance to unload the sand particle. If a termite arrives at the tunnel face to find another termite already excavating there, then it chooses between waiting for the current excavator to finish or instead excavating the sidewall and thus starting a new branch tunnel (Figs. 4B, S2). We used the species-specific probabilities of these behaviors in the simulations, based on the experimental results for *R. tibialis* and *H. aureus* (Fig. 4AB). We also simulated building by *P. simplicicornis* to predict if the same mechanism can explain the branching patterns of this species. For them, we added the behavior that one termite can take over sand particles from another termite. In the simulation, the side length of a single cell is 10 mm, and we observed the development of tunnels until the longest path reached 100 mm. As in the experiments, we characterized branching pattern by counting the number of tunnel faces.

The model effectively reproduced the interspecific variation among rhinotermitid termites, where *R. tibialis* built tunnels with less branching than *H. aureus* (Fig. 4C). However, the model underestimated the branching rates of *P. simplicicornis*, suggesting that this species may have another behavioral mechanism for branching in addition to sidewall excavation within a clogged tunnel. Moreover, our model revealed that high local density causes branching in termite tunnels. Our experiments showed that the branching of termite tunnels is concentrated near the beginning of tunnels (Fig. 2D). The same pattern was reproduced by our simulations (Fig. S3). This is because, at the beginning of the excavation, the area of the tunnels is smaller, which increases the local density of termites and the probability of sidewall excavations. As the tunnels grow, the area increases and local density declines, which results in lower branching rates in later stages of excavation. It is well known that the group size and density affect the tunneling structures in many social insects [26–28]. But our results indicate that even with the same group size, the change of the local density of individuals will greatly affect pattern formation.

## Discussion

Our comparative study revealed a complex relationship between behavioral mechanism and group-level patterns. We found that two closely related species (*R. tibialis* and *H. aureus*) share behavioral repertoires, but quantitative differences in the frequency of different actions result in divergent branching patterns (Fig. 4). This result shows that parameter tuning of the same rule set plays an important role in the evolution of collective building, and thus that a dramatic change of behavioral repertoires is not required to produce diverse nest structures among species. In contrast, we also found that two phylogenetically divergent species (*P. simplicicornis* and *R. tibialis*) possess different behavioral repertories for collective excavation, but this does not necessarily yield different structures, since both create tunnels with a similar branching pattern (Figs 2, 3). Thus, similarity of patterns need not imply a shared behavioral algorithm. Altogether, we conclude that the evolutionary process of collective behavior is much more complex than the transition of group-level patterns. This makes it impossible to solve the inverse problem of inferring individual behavioral rules from the collective structures that they produce. Our result emphasizes the importance of direct comparative studies of behavioral mechanisms of self-organizing systems.

Termite phylogeny shows that tunneling behavior was present in the common ancestor of Rhinotermitidae (Fig. 1). This suggests that parameter tuning of shared behavioral repertories explains pattern diversification in this whole group. The Rhinotermitidae have a wide diversity of tunneling patterns [19,20], which are often connected to different foraging strategies [21,29,30]. Optimal search theory predicts a compromise between reducing detection errors and quickly exploring a wide area [31]. In this sense, *H. aureus* engages in an intensive search by building highly branched tunnels, while *R. tibialis* performs an extensive search by focusing on fewer tunnels. This may reflect their habitats, where *H. aureus* is found in deserts with cactus resources that are small and relatively difficult to find, while *R. tibialis* lives in pine forest with large wood resources. However, other factors including colony size and traveling cost also determine search efficiency [32,33]. We found that the behavioral mechanism underlying this adaptation is simple, where the tunnel geometry is sensitive to a single behavioral parameter governing interactions; namely, a threshold for individuals in a clogged tunnel to excavate a sidewall instead of waiting for access to the tunnel face. Thus, without the need for a change in communication systems, selection acting on this parameter can easily result in the evolutionary divergence of tunnel geometry, adapting it to the local environments experienced by each species. This tunability of behavioral algorithm may have helped the Rhinotermitidae to spread to a wide range of environments [34].

On the other hand, the evolution of a differentiated behavioral rule in *P. simplicicornis* indicates the importance of evolutionary contingency. This species’ behavioral repertoire appears to have been shaped by the physiological and morphological traits of its family, Kalotermitidae. First, kalotermitids move slower than rhinotermitids, possibly because of lower metabolic rates or shorter legs [35,36]. During our observations, the maximum instantaneous moving speed of *P. simplicicornis* was significantly slower than that of *R. tibialis* or *H. aureus* (Fig. S4). Kicking works well for slower moving termites because it requires a shorter total movement distance to excavate a unit length of tunnel (Fig. S4). Second, the body shape of kalotermitids is more elongated than that of rhinotermitids (Fig. S5), which limits turning around inside narrow tunnels [37]. Because of this characteristic, *P. simplicicornis* may do better with the kicking type of tunneling. Indeed, turning behavior, which often involves transportation of sand particles for a longer distance, is less frequently observed in *P. simplicicornis* (Fig. S5). Thus, the kicking type of tunneling is an adaptation to confined space for species with lower mobility.

When a group of animals moves within a narrow and confined space, they face a problem of high-density clogs which affect task performance and collective outcomes [27,37]. The bucket brigade is one solution, because excavators do not need to pass each other [38]. In addition to *P. simplicicornis*, there are a few observations of this behavior in social mole-rats [39] and army ants [40]. Another mechanism, observed in fire ants, is individual idleness, which limits the number of excavators at the face of tunnels and reduces the frequency of clogs [37]. The higher proportion of waiting behaviors by *R. tibialis* is consistent with this idea. Thus, there are different clog control mechanisms behind the convergence of reduced tunnel branching in *P. simplicicornis* and *R. tibialis*. Instead of reducing local density, *H. aureus* exploits high-density clogs as a mechanism for building new branches, thus forming highly branched tunneling patterns. A similar mechanism of high density creating a new branch has also been proposed for ant nest construction [27]. Combined with previous studies, our results illustrate that collective behavior in confined space is flexible within each animal taxon, suggesting that each group’s behavior is an adaptive trait.

The central goal of evolutionary biology is to find the factors and mechanisms which are responsible for the evolution of phenotypes. Two striking evolutionary phenomena are divergence of phenotypes among closely related species and convergence between distinct related species. Studies on morphological development of individual animals have found that there is not a one-to-one relationship between morphology and its genetic determinants; different forms can evolve by altering the expression of shared genes [17], and similar forms can evolve by using a different molecular basis [16]. Under the superorganism analogy, the nest is the extended phenotype of an insect colony, formed by the behavioral rules of colony members. Consistent with studies on morphological development, our study reveals that diverse group-level patterns between species are produced from shared behavioral rules, and also provides a novel example of convergent extended phenotypes based on distinct behaviors. This challenges theoretical studies that often assume the same behavioral rules across taxa based on the observation of a limited number of species. Direct comparative studies promise a comprehensive view of the mechanisms of collective behavior and will give us an understanding of the origin of coordination, as well as the general algorithms underlying it.

## Materials and Methods

### Termites

We used four colonies of *Paraneotermes simplicicornis* (Kalotermitdae) and five of *Heterotermes aureus* (Rhinotermitidae) collected from cholla and mesquite desert in Gila and Maricopa Counties, and five colonies of *Reticulitermes tibialis* (Rhinotermitidae) collected from a pine forest in Pinal County, Arizona, USA. *P. simplicicornis* is the sole subterranean termite species in Kalotermitidae [41], while all species in *Heterotermes* and *Reticulitermes* are counted as subterranean termites. Colonies were maintained at 22 °C in plastic boxes with wood or cactus pieces and the soil in which they were nesting in the field.

### Macro-scale observation of tunneling patterns

To compare the branching patterns of tunnels, we prepared experimental setups for observing two-dimensional tunneling patterns. These consisted of three layers. The middle layer, whose thickness is adjusted to each species (1 mm for *H. aureus* and *R. tibialis* and 2 mm for *P. simplicicornis*) had a round area filled with white sand (Marble White Sand, National Geographic, USA) moistened with distilled water (10% by volume). At the edge of the round area was a teardrop-shaped area where termites could be introduced (Fig. 2A). Sand particles were homogenized into 0.15 ∼ 0.25 mm size range using two screens with 60 and 100 mesh. The top layer had an opening only above the entry area, which was covered by a glass plate. We used 20 termites for this experiment. Each individual was used only once. After placing termites in the introduction area, we recorded tunnel development for 24 hours. Snapshots were imported into ImageJ (US National Institutes of Health, Bethesda, MD, USA) and measurements were taken by tracing the length of each branch after calibration. We defined the beginning of the tunnel structure as a single point connected to the entry area. Tunnels greater than one body length (6.2 mm, 4.4 mm and 3.9 mm for *P. simplicicornis, R. tibialis*, and *H. aureus*, respectively) were counted as unique branches.

Overall, *P. simplicicornis* formed tunnels much more slowly than *R. tibialis* and *H. aureus* (Fig. S6). To avoid an effect of environmental heterogeneity arising from the wall at the boundary [42], we compared the structures of tunnels at the time when the first group in each species reached the wall (14 hours in *P. simplicicornis* and 5 hours in *R. tibialis* and *H. aureus*; Fig. S6). We compared the number of tunnel faces among species using a generalized linear model (GLM) with Poisson error and a log-link function. The likelihood-ratio test was used to test for statistical significance of the explanatory variable (type-II test). We pooled the data of three colonies for each species because we did not find any significant colony variation (GLM, likelihood-ratio test, *P* > 0.12). In case of significant effect of species, we ran Tukey’s post hoc test.

### Micro-scale observation of digging behaviors

To compare micro-scale digging behaviors among the three species included in our study, we prepared experimental arenas of two different sizes depending on species (small: *H. aureus* and *R tibialis*; large: *P. simplicicornis*). The experimental arenas consisted of three layers; bottom and middle layers made of acrylic board and a top layer made of glass plates. The middle layer was L shaped and included an empty square area for introducing termites (small: 15 × 15 × 1 mm; large: 20 × 20 × 2 mm) and a narrow passage (small: 3 × 100 × 1 mm; large: 4 × 100 × 2 mm) filled with white sand (Fig. S7A).

We selected ten termites haphazardly from available colonies for each trial of this experiment. We used workers for *H. aureus* and *R. tibialis*; for *P. simplicicornis* we used pseudergates or nymphs who play the role of the worker caste in Kalotermitidae, which have no true worker caste [43]. Each *P. simplicicornis* group contained either all pseudergates or all nymphs. All termites were marked with one dot on the head and two dots on the gaster (Racing Finish, Pactra, Testors, Rockford, IL, USA). We used the marking on the head for tracking, and those on the gaster for individual identification. After installing termites in the square area, we recorded their behaviors until they dug a tunnel 50 mm long; as in the two-dimensional experiment, *P. simplicicornis* took longer to reach this milestone (Fig. S6). A video camera was mounted above the arena covering the square area and first 50 mm of the passage. We observed three groups from two colonies for each species; one replicate for *P. simplicicornis* (colony A, rep 3) was censored at 24 hours after introduction of the termites, when the tunnel had reached 47.60 mm in length. We used each individual only once.

All videos were split into 30 minute segments. Then we identified the segment in which the termites started excavation. Starting with the immediately following segment, we observed their behavior for 10 minutes every 60 minutes. During observations, we extracted the coordinates of each termite’s head at a rate of 1 frame per second from each video using the video-tracking system UMATracker [44]. We also measured the length of the tunnel at the beginning and the end of each observation.

By analyzing the trajectories, we obtained the number of visits to the tunnel face by 1st-row individuals and the number of changes in position between the 1st- and the 2nd-row individuals. We estimated the mean numbers of these behaviors performed during the digging of a 1mm length of tunnel. Then we compared the mean frequency of these behaviors among species using one-way analysis of variance (ANOVA). We pooled the data of two different colonies for each species because we did not find significant colony variations for any species (t-test, *P* > 0.10).

To form the tunnel, termites visit the tunnel face, excavate sand, and then transport sand particles away from the tunnel face. Because of the narrow tunnel, only the 1st-row individuals can access the tunnel face. Thus, we focused detailed analysis on the behavior of 1st-row individuals, and we determined their behavioral repertories when the tunnel is longer than 40mm. We considered the 1st-row individual to have visited the tunnel face when its position came within 1.5mm of the tunnel face and then backed away more than 2mm (or 3mm for *P. simplicicornis*). We observed these visits to check if the termites excavated sand, how they carried sand particles, and where and how they deposited them. Next, we examined the interaction patterns among individuals by focusing on the 2nd-row individuals who are found just behind a 1st-row individual visiting the tunnel face. We only considered 2nd-row individuals that were within a minimum distance of the 1st-row termite. This minimum distance was 6.5 mm, 4.5 mm and 4 mm for *P. simplicicornis, R. tibialis*, and *H. aureus*, respectively (i.e., a little larger than body length for each species). The frequency of observed behaviors was compared among species using Fisher’s exact test.

### Individual-based model

We modeled a 2D discrete space, representing positions in tunnels. Each cell had two possible states (empty and sand-filled), and termites were modeled as mobile agents, each one occupying a single empty cell. All termites were initially placed in the introduction area, which can contain all individuals.

Termites have five different states: moving forward (advancing), excavating, backing (with or without loading) and waiting. Inside a tunnel, termites determine their behaviors depending on their state and their interactions with other individuals. Each behavior takes one time step (Fig. S2). If individuals don’t encounter others, advancing termites move forward as long as the cell in front is empty. When the front is sand filled, advancing termites change to the excavating state; then they excavate sand and change to the state of backing while loaded with sand. Backing termites move back as long as the cell behind them is empty, until they have moved a given backing distance or left the tunnel. As the backing distance increased proportionally to the tunnel length (Fig 3D), we determined the backing distance by multiplying the tunnel length by a random number generated from a beta distribution with parameters α and β (Table S1), which are variable among species and obtained by fitting to observed transport distances (data are shown in Fig. S5). We used a beta distribution because it can describe the bimodal shape observed in the data. After backing, individuals unload the sand and enter the advancing state. Once unloaded, the sand is no longer available in simulations of *R. tibialis* and *H. aureus*, but remains available in *P. simplicicornis*, where other individuals can pick it up in a time step.

When there is an excavating individual in front of an advancing termite, it will enter either the waiting or excavating state, with probabilities based on those observed for waiting and sidewall excavation in the experiments (Fig. 4A). Then, waiting individuals will swap positions with the individual in front once its state changes to backing; excavating individuals will excavate the sidewall to create a new branch. When there is a backing individual in front of an advancing termite, the latter changes to the backing state because of the confined space. Moreover, in *P. simplicicornis*, when an advancing individual encounters a backing individual who is loaded with sand, the advancing termite takes over the sand particles. After this, the backing individual changes to advancing state, while the advancing individual changes to backing state. Similarly, individuals of *P. simplicicornis* can also choose to take over a sand load when they encounter excavating individuals (Fig. S2).

In each trial, we modeled 20 individuals, as in the experiments. These 20 individuals act sequentially in random order at each time step, except for swapping or taking over where we need to compute the action of two individuals at the same time. The side length of a single cell is 10 mm, and we observed tunnel development until the longest path reached 100 mm. As in the experiments, we ran the simulation 15 times for *R. tibialis* and *H. aureus* and 16 times for *P. simplicicornis*. Then we counted the number of tunnel faces and measured the mean values. We repeated this process 1000 times to estimate the expected range of the mean value. All calculations were performed using R v.3.5.3 [45].

## Acknowledgments

We thank K. Baudier and C. Kwapich for helpful comments during analysis; T. Bourguignon for information on termite phylogeny; M. De La Monja and A. Rizo for help during image analysis; members of the Pratt laboratory and the Social Insect Research Group at ASU for helpful discussion. N.M. is supported by a JSPS Overseas Research Fellowship.

## Supplemental materials

### SI text

#### Phylogenetical analysis

To reconstruct the evolutionary of tunneling behaviors of lower termites, we first mapped the presence of tunneling behaviors on the phylogeny. We assumed that the multiple-piece or central nesting termites do tunneling through the soil, while single-piece nesting termites do not [46]. Although there are some observations of tunneling behaviors even in single-piece nesting termites [47–51], the frequency of tunneling behaviors in the former is much higher than the later. Moreover, note that the wood roach *Cryptocercus*, which is the sister group of all modern termite species, does not dig in soil but excavates inside the gallery of well-rotten logs [52]. Because of the lacking of the study, the information of Rhinotermitidae is limited. In this study, we follow the analysis in Inward et al. 2007, to assume that only *Prorhinotermes* is classified as a single-piece nester [53]. Recent molecular phylogeny can be found in Bourguignon et al. 2015 [24], but this does not include genus *Paraneotermes*. Thus, based on the morphological phylogeny [23], we inserted *Paraneotermes* and *Kalotermes* into the molecular phylogeny. We conducted the ancestral state estimates using a maximum parsimony approach implemented in Mesquite v3.6 [54].

**Figure S1.**
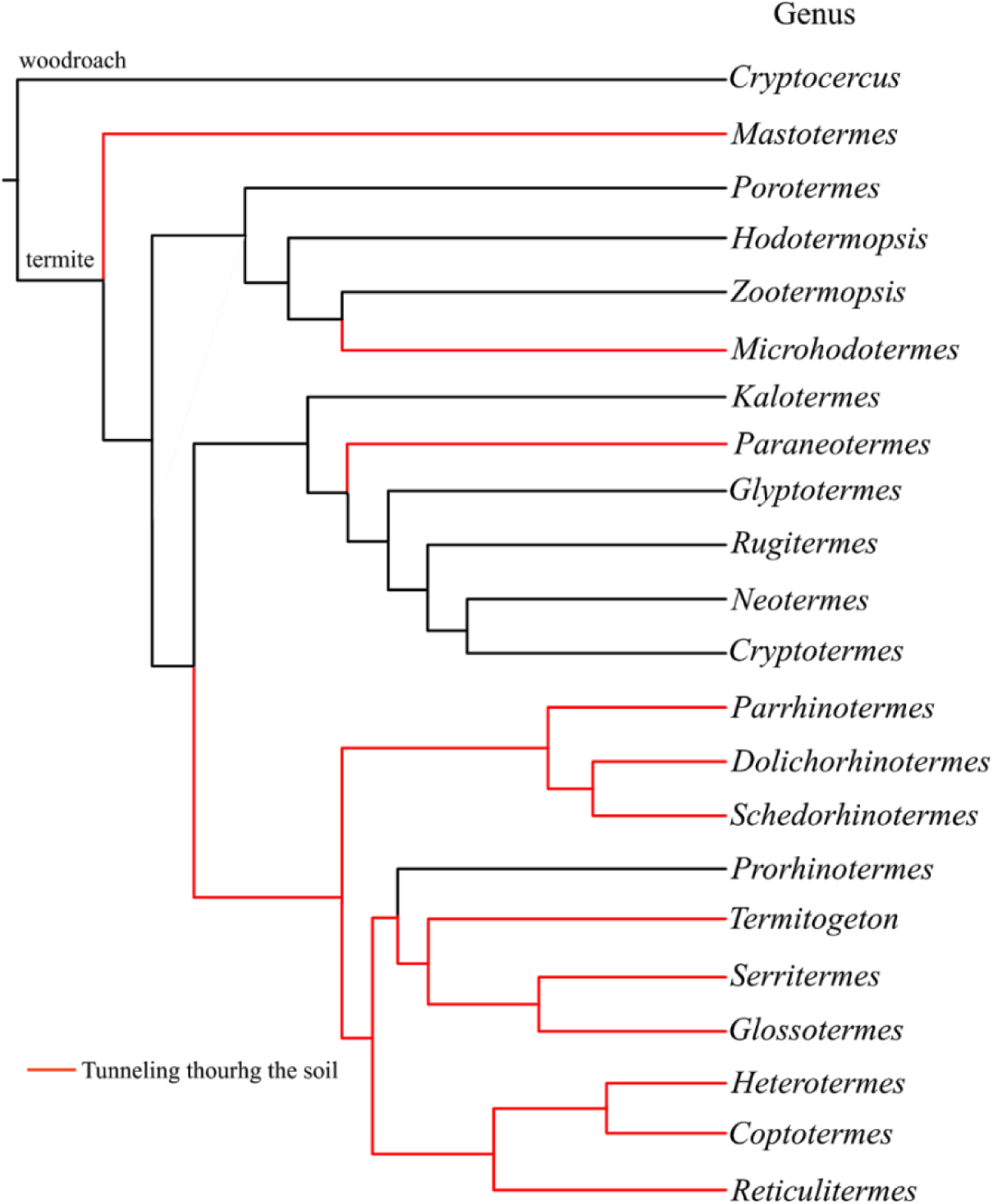
Detailed phylogeny of lower termites (modified from [23,24]) with the information of tunneling behaviors. The ancestral states were reconstructed with maximum parsimony. Tunneling through the soil has evolved four times independently in Mastotermitidae, Hodotermitidae, *Paraneotermes*, and Rhinotermitidae.

**Figure S2.**
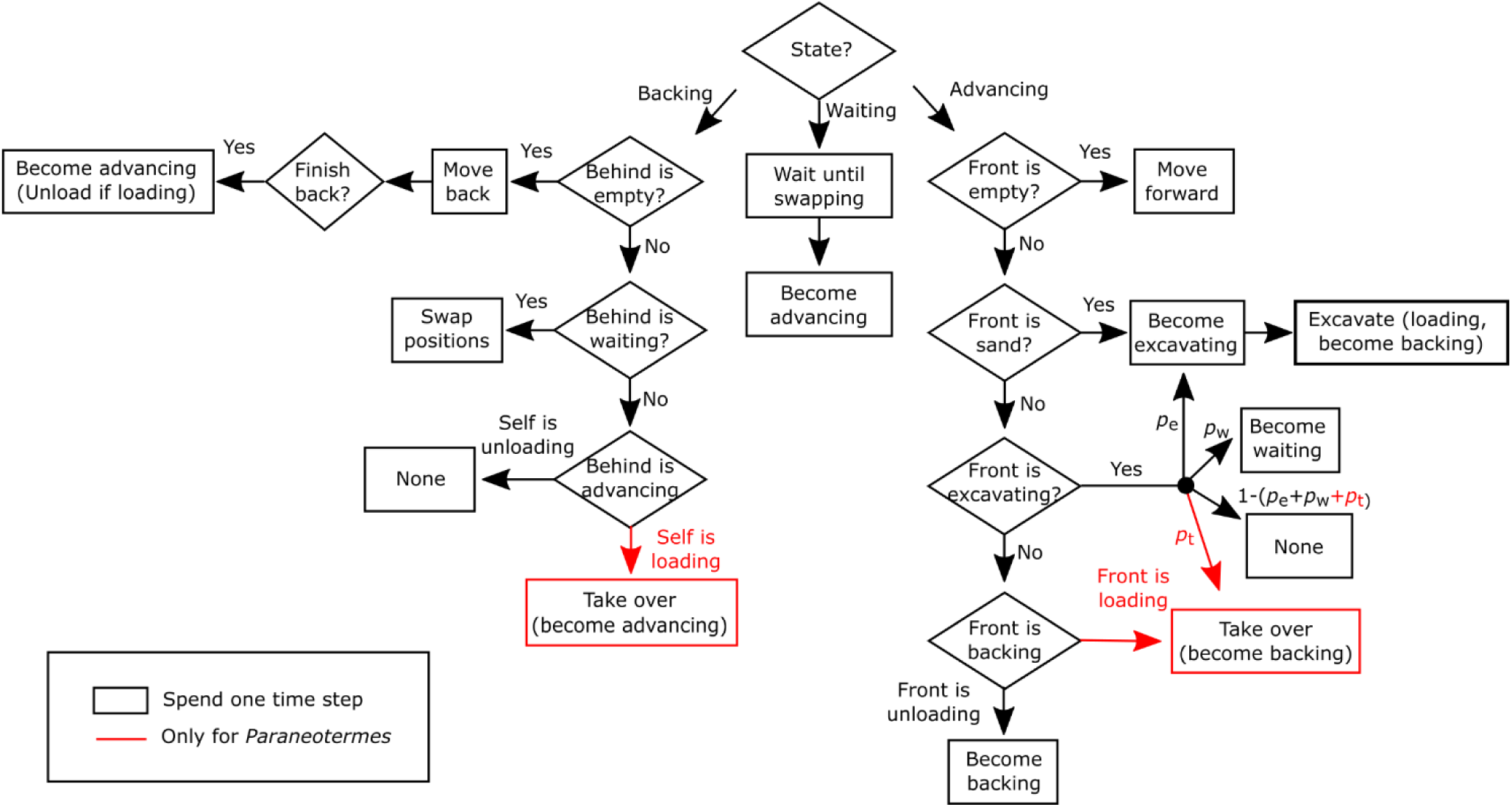
Model of termite collective excavation. The flowchart begins at the top with “State?”. Once it reaches the endpoint, it goes back to the beginning, “State?”. Parameter values of *p*_e_, *p*_w,_ and *p*_t_ are in Table S1.

**Figure S3.**
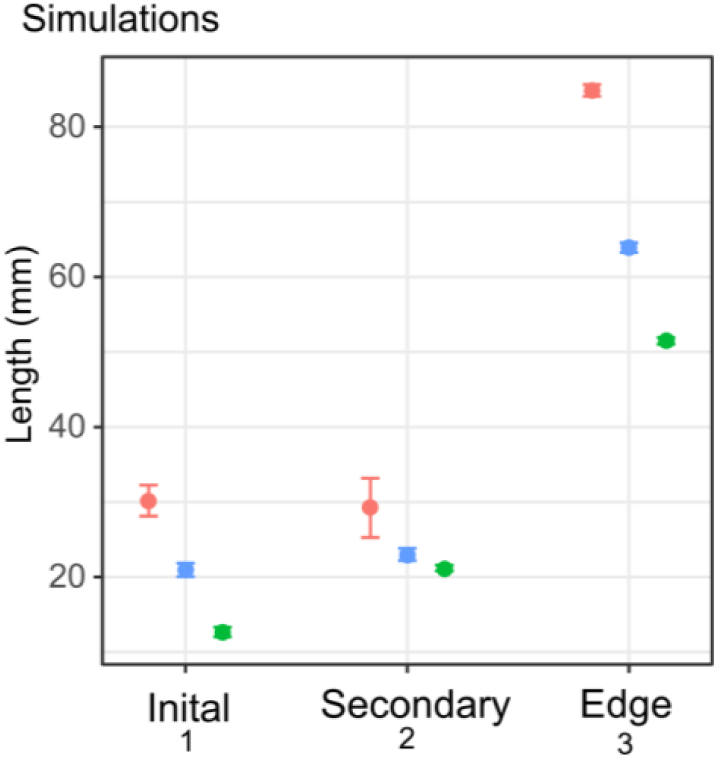
Comparison of the tunnel length for the simulations when divided into segments. Initial tunnel is the segment from the entrance to the first branch. Secondary tunnel is a segment between two branches. Edge tunnel is a segment reaching the face of a tunnel. When a tunnel has no branch, it only contains an edge tunnel. See also Fig. 2D.

**Figure S4.**
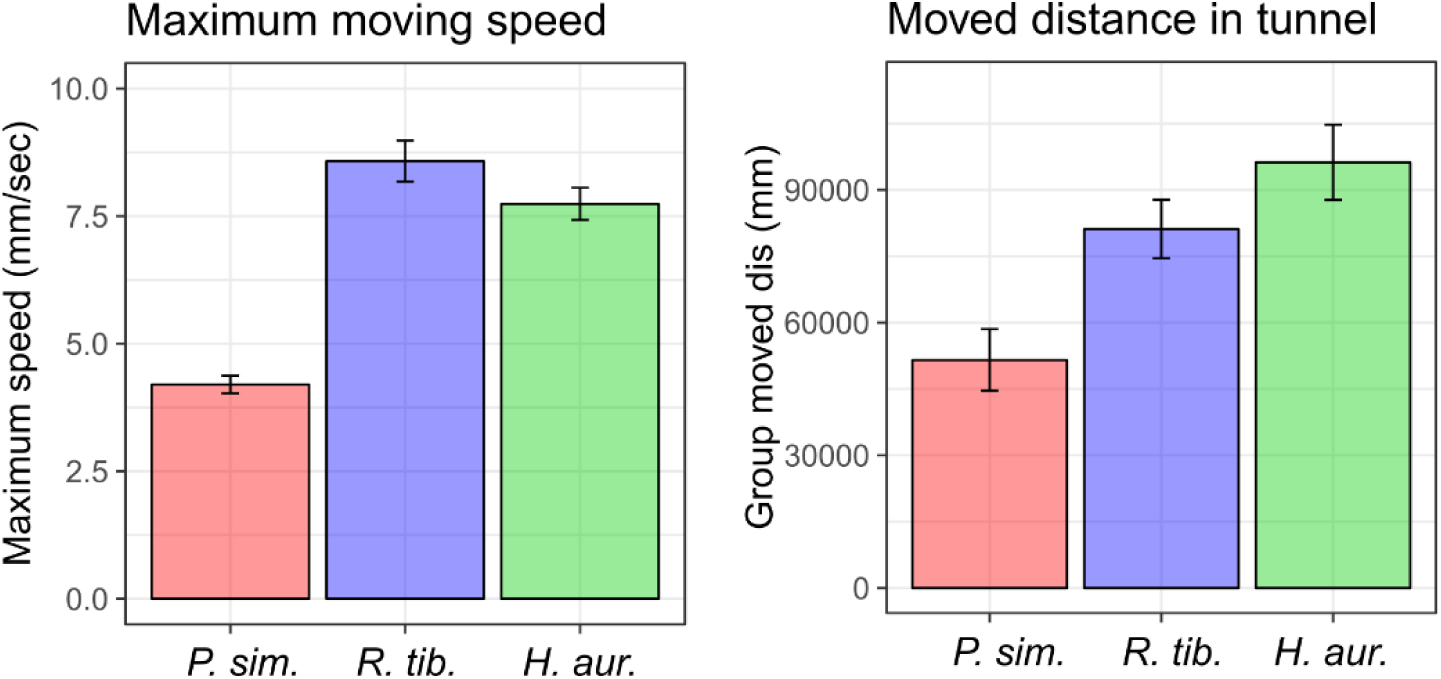
Comparison of the maximum instantaneous moving speed and total moved distance during observations. Maximum moving speed was measured for each individual separately, while moved distance was measured for each group by summing up the distance traveled by all members in a group inside a tunnel. As the observation was performed for 10 minutes every hour, we multiplied the distance by 6 to get the estimated values. Bars indicate mean ± S.E.

**Figure S5.**
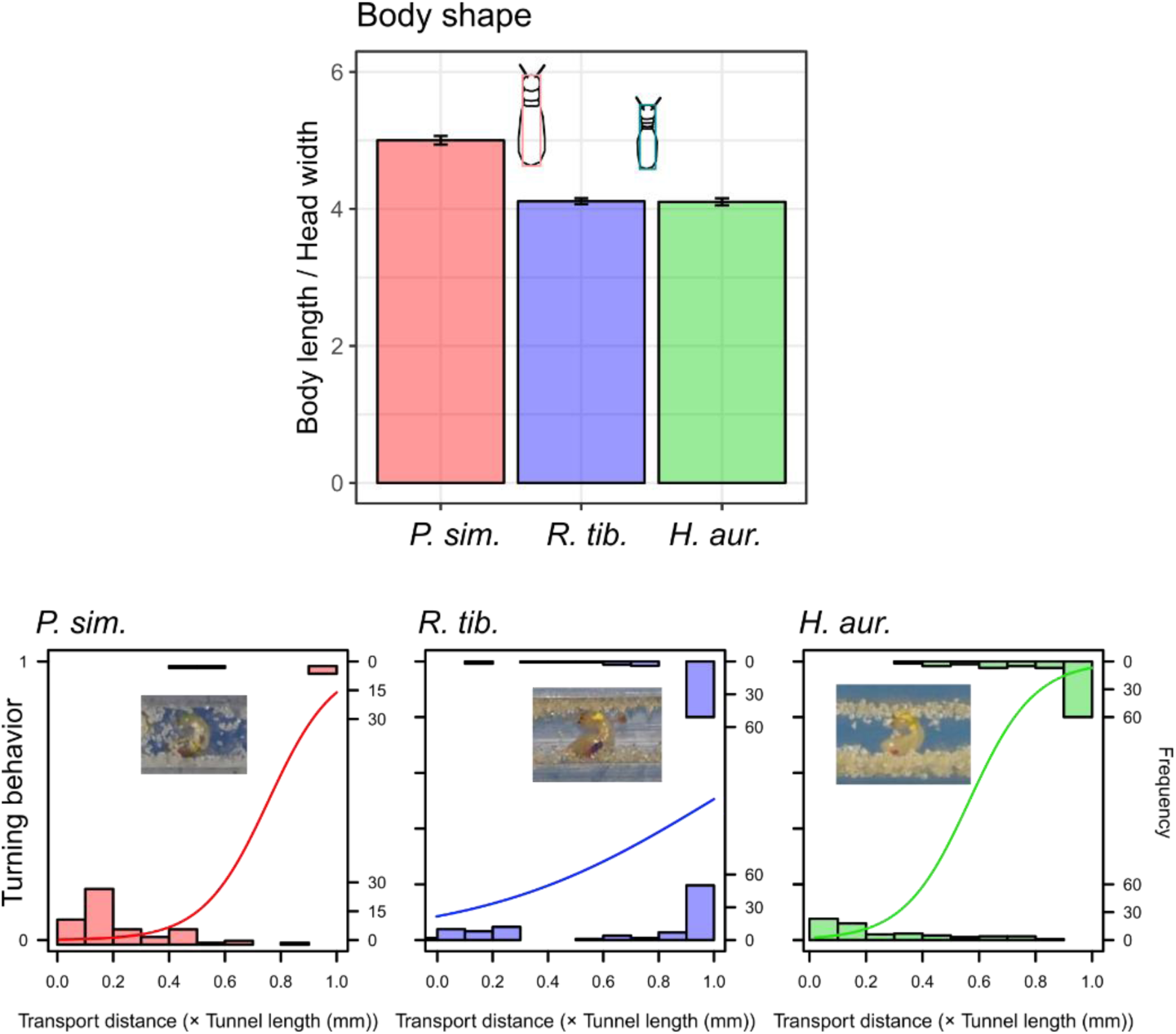
Comparison of body shape and turning behavior inside a tunnel. Body length and head width were measured for 30 inidviduals for each species. Bars indicate mean ± S.E. Turning behaviors were measured for 1st row individuals which leave the tunnel face after excavate and when the length of the tunnel is longer than 40mm. Logistic regression curve is shown for each species, where we tested the relationship between transport distance and presence of turning behavior.

**Figure S6.**
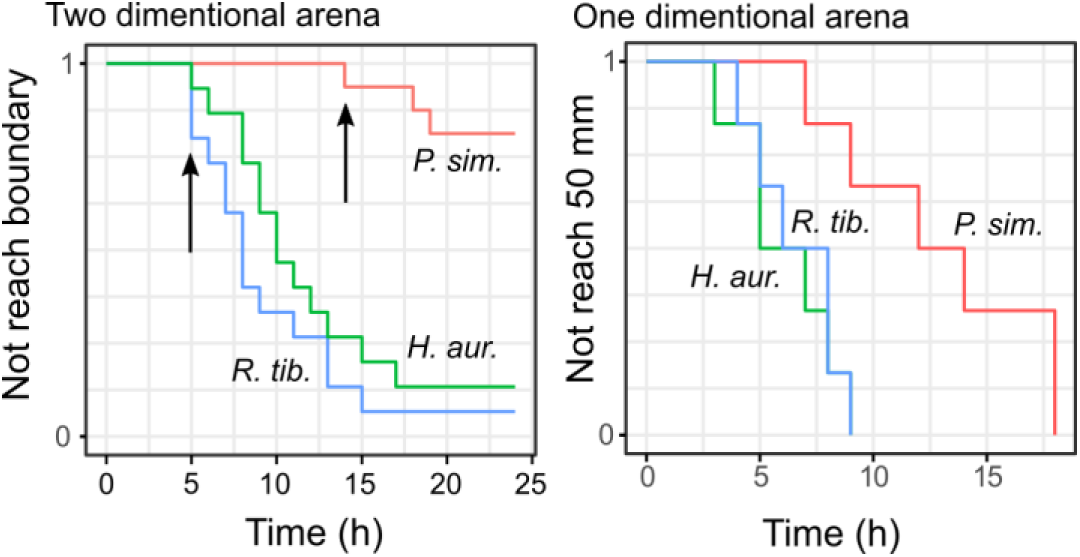
The time until a group of termites reach a wall (two dimensional) or create a 50mm tunnel (one-dimensional arena). Arrows indicate the timing we analyzed the tunneling patterns (two dimensional).

**Figure S7.**
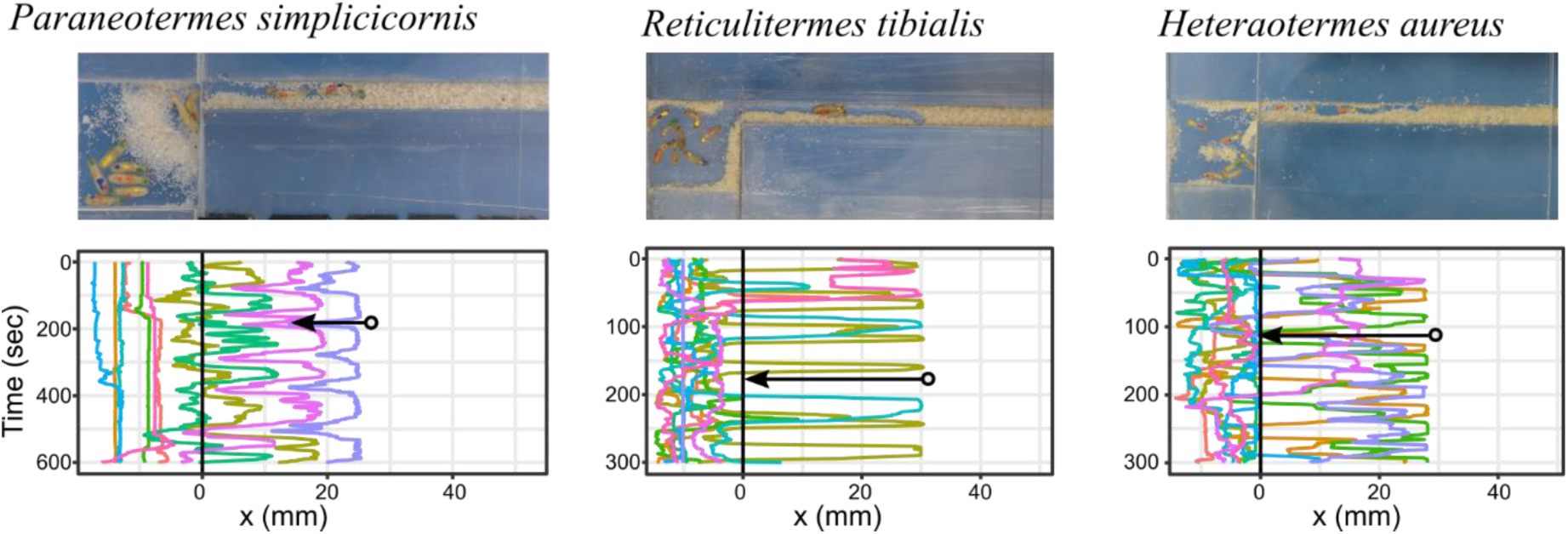
Experimental setup for behavioral observations with representative x-axis trajectories of each termite. Different individuals are shown in a distinct color. Black arrows show examples of backward movements, where *R. tibialis* and *H. aureus* often go back to the entrance of the tunnel, while *P. simplicicornis* mainly move only within a limited range within the tunnel.

**Table S1.**
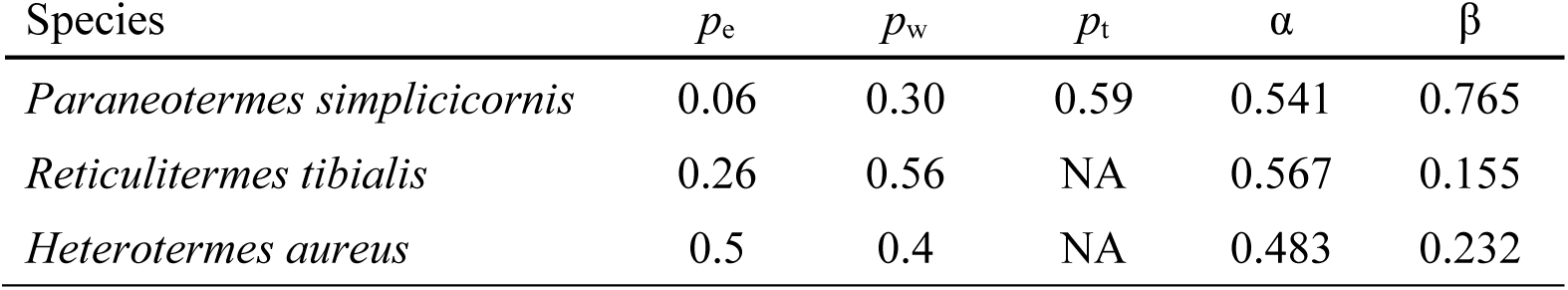
Parameter values used in our simulations.

